# Survival of *Campylobacter jejuni* in Amoebae enhances subsequent invasion of mammalian cells

**DOI:** 10.1101/2021.11.25.469992

**Authors:** Fauzy Nasher, Brendan W. Wren

## Abstract

The ubiquitous unicellular eukaryote, *Acanthamoeba*, is known to play a role in the survival and dissemination of *Campylobacter jejuni. C. jejuni* is the leading cause of bacterial foodborne gastroenteritis world-wide and is a major public health problem. The ability of *C. jejuni* to interact and potentially invade epithelial cells is thought to be key for disease development in humans. We examined *C. jejuni* grown under standard laboratory conditions,11168H_CBA_ with that harvested from within *Acanthamoeba castellanii* (11168H_AC/CBA_) or *Acanthamoeba polyphaga* (11168H_AP/CBA_), and compared their ability to invade different cell lines. *C. jejuni* harvested from within amoebae had a ∼3.7-fold increase in invasiveness into T84 human epithelial cells and a striking ∼11-fold increase for re-entry into *A. castellanii* cells. We also investigated the invasiveness and survivability of six diverse representative *C. jejuni* strains within *Acanthamoeba spp*., our results confirm that invasion and survivability is likely host cell dependent. Our survival assay data led us to conclude that *Acanthamoeba spp*. are a transient host for *C. jejuni* and that survival within amoebae pre-adapts *C. jejuni* and enhances subsequent cell invasion. This study provides new insight into *C. jejuni* interactions with amoebae and its increased invasiveness potential in mammalian hosts.

## Introduction

*Campylobacter jejuni* is the leading cause of bacterial food-borne gastroenteritis worldwide [1]. However, it is puzzling that this microaerophile bacterium that is incapable of growing under atmospheric conditions [2] can be so prevalent in the environment and being responsible for such widespread disease in humans. It is still unclear how this pathogen survives and thrives in the environment outside its warm-blooded avian and mammalian hosts. Several studies have reported survival of *C. jejuni* within free living protozoa, such as amoebae, as a mode of survival and persistence in the environment [3-5].

Free-living amoebae are widely distributed in the environment and have been isolated from a range of sources including freshwater, seawater, soil, dust and food sources [6, 7]. Amoebae, including *Acanthamoeba spp*., have long been investigated for their role to phagocytose bacteria as prey, to serve as a vector or a host to pathogenic bacteria, including *Campylobacter spp*., *Legionella spp*., *Mycobacterium spp*. and *Pseudomonas spp*. [5, 8, 9]. *Acanthamoeba spp*. as a vector and/or a host include aiding in bacterial survival with or without multiplication. Growth and multiplication of bacteria can lead to subsequent lysis of the amoebae and release of bacteria [9]. This “Trojan horse” principal for bacterial pathogens has been linked to disease outbreaks in contaminated water [10] and food sources [11].

The phenomenon of *C*.□*jejuni* survival within amoebae has been previously studied without any definitive insight into its role in human disease. Here, we show that survival within amoebae pre-adapts *C. jejuni* and enhances subsequent invasion to mammalian cells which could lead to increased disease in mammalian hosts including humans.

## Materials and Methods

### Strains and Cultures

Bacteria were stored using Protect bacterial preservers (Technical Service Consultants, Heywood, U.K.) at −□80 °C. *C. jejuni* strains were streaked on blood agar (BA) plates containing Columbia agar base (Oxoid) supplemented with 7% (v/v) horse blood (TCS Microbiology, UK) and Campylobacter Selective Supplement (Oxoid), and grown at 37 °C in a microaerobic chamber (Don Whitley Scientific, UK), containing 85% N_2_, 10% CO_2_, and 5% O_2_ for 48 hrs. *C. jejuni* strains were grown on CBA plates for a further 16 hrs at 37 °C prior to use.

*Acanthamoeba castellanii* CCAP 1501/10 and *Acanthamoeba polyphaga* CCAP 1501/14 (Culture collection of Algae and protozoa (Scottish Marine Institute)) were grown to confluence at 25°C in 75-cm^2^ tissue culture flasks containing peptone yeast and glucose (PYG) media [12]. Amoebae were harvested by scraping the cells into suspension, and viability was determined by staining with trypan blue and counting by a hemocytometer using light microscopy.

### *C. jejuni* invasion and survival assay

*C. jejuni* 11168H, a derivative of the original sequence strain NCTC 11168 was used in this study. *C. jejuni* cells were either grown on CBA agar (11168H_CBA_) as described above or harvested after intracellular survival in *A. castellanii* (11168H_AC/CBA_) or *A. polyphaga* (11168H_AP/CBA_) **(Table 1)**, before invasion of epithelial cells and re-invasion of amoebae. Briefly, a large-scale invasion of *Acanthamoeba spp*. was carried out in a 150 cm^2^ tissue culture flask (Falcon), a monolayer of approximately 10^6^ amoebae was infected with *C. jejuni* at a multiplicity of infection (MOI) of 200:1 for 3 hrs at 25 °C in PYG media. The monolayer was washed 3x with 25 ml of PYG media and incubated for 2 hrs in 25 ml of PYG media containing 100 μg/mL of gentamicin. *C. jejuni* cells were harvested by scraping the amoebae into suspension and centrifuged for 10 min at 350 x g to pellet the bacteria and amoebae. Supernatant was discarded and the pellet was suspended in 1 ml of distilled water containing 0.1% (v/v) Triton X-100 for 10 min at room temperature to lyse the amoebae and release bacteria cells. The suspension was then centrifuged for a further 10 min at 4000 x g, the resultant pellet was resuspended in 1 mL PBS and 200 μl aliquots of this suspension were plated on CBA plates and incubated microaerobically for 48 hrs at 37 °C to ensure recovery of enough bacteria. For invasion assay, the experiment was performed as described above, with an additional step of serial dilutions and plating out bacteria for colony forming units (cfu). *C. jejuni* invasion of *Acanthamoeba spp*. was confirmed by laser scanning microscopy (LSM) after 3 hrs infection with 11168H_GFP_ *C. jejuni* strain (**Fig. 1**).

**Table 1.** Representative *Campylobacter jejuni* strains used in this study.

| STRAIN | DESCRIPTION | MULTI LOCUS<br>SEQUENCE TYPE | REFERENCE |
| --- | --- | --- | --- |
| **11168H** | A hyper-motile derivative of the original sequence strain NCTC 11168 that shows higher levels of caecal colonization in a chick colonization model | ST-21 | Karlyshev *et al.*, [36]<br>Jones *et al.*, [37] |
| **81-176** | Highly virulent and widely studied laboratory strain of *C. jejuni*. MLST | ST-42 | Korlath *et al.*, [38] |
| **12912** | Ox liver portion isolate | HS Type-50 | Gundogdu *et al.*, [39] |
| **M1** | A rarely documented case of direct transmission of *C. jejuni* from chicken to a person, resulting in enteritis | ST-45 | Friis *et al.*, [40] |
| **81116** | Genetically stable strain which remains infective in avian models | ST-283 | Wassenaar *et al.*, [41] |
| **RM1221** | A chicken isolate with unique lipooligosaccharide and ability to colonize chicken skin | ST-354 | Fouts *et al.*, [42] |
| **11168H<sub>CBA</sub>** | Grown under standard laboratory conditions |  | This study |
| **11168H<sub>AC/CBA</sub>** | Strain 11168H harvested after survival in *Acanthamoeba castellanii* |  | This study |
| **11168H<sub>AP/CBA</sub>** | Strain 11168H harvested after survival in *Acanthamoeba polyphaga* |  | This study |
| **11168H<sub>GFP</sub>** | Strain expressing green fluorescent protein |  | Jervis *et al.*, [43] |

**Figure 1.**
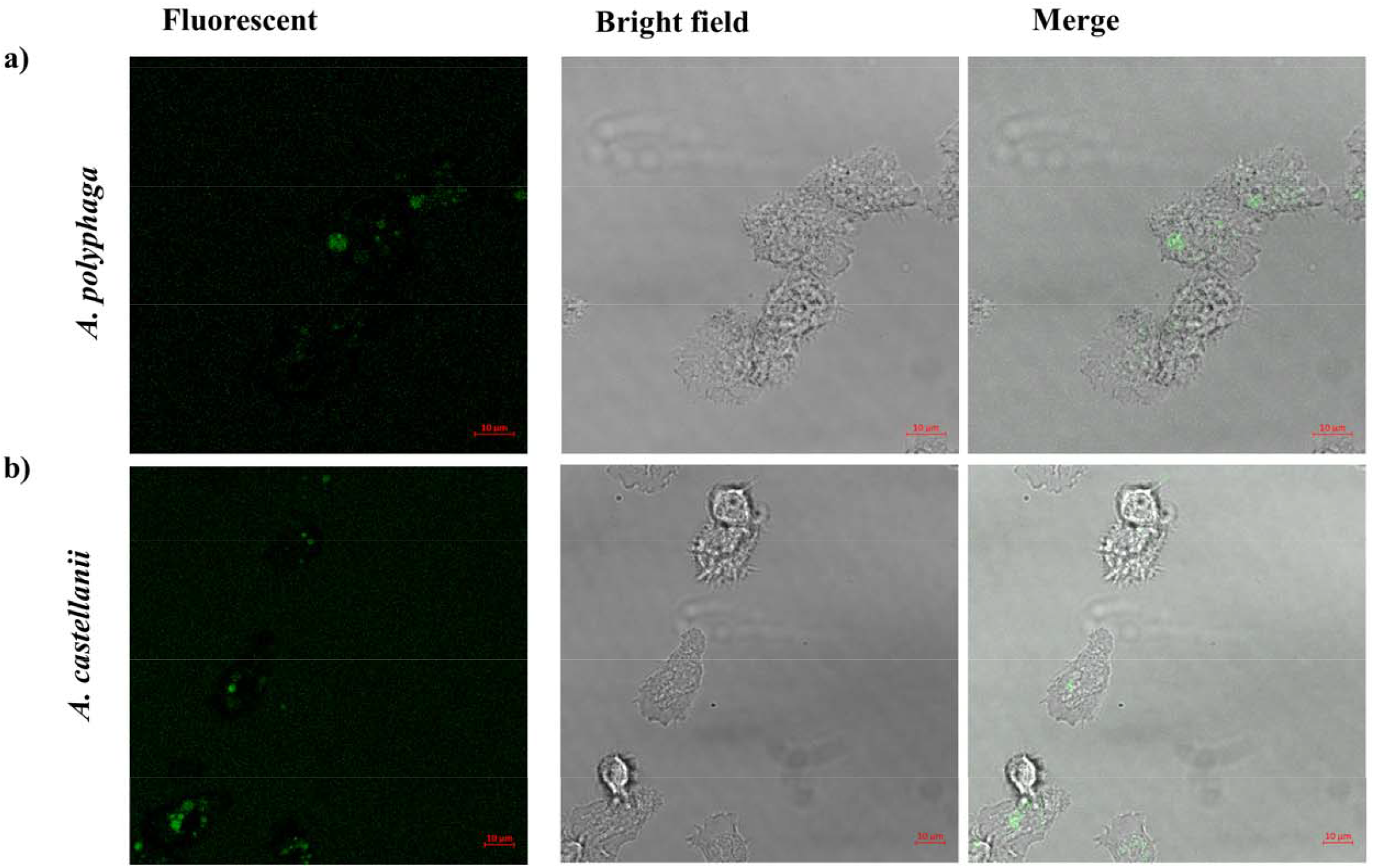
Laser scanning confocal microscopy (LSM). Observation of GFP labelled *C. jejuni* strain 11168_GFP_ within **a)** *A. polyphaga* and **b)** *A. castellanii* after 3 hrs infection. *C. jejuni* is observed as green fluorescent within the amoebae cells. A monolayer of amoebae (10^6^) in a 35 mm imaging dish (Ibidi) were infected with *C. jejuni* to an M.O.I of 200 and incubated for 3 hrs aerobically at 25 °C, cells were washed 3x before imaging. Laser scanning microscopy (Axion) was used to image the cells at objective 63x/1.4 oil. *C. jejuni* 11168_GFP_ strain was constructed as described previously[43].

For survival assay, the experiment was performed as described above with the following modifications; an additional step of incubation with 10 μg/mL of gentamicin was added before the cells were washed at each timepoint and lysed to plate for enumeration as described above.

### Human cell lines and culture conditions

T84 (human carcinoma cell line) and Caco-2 (human colorectal adenocarcinoma cells) were grown in Dulbecco’s modified Eagle’s medium and Ham’s F-12 (DMEM/F-12) supplemented with 10% fetal bovine serum (FBS) and 1% non-essential amino acid. The monolayers, ∼10^5^, were seeded in a 24-well tissue culture plates and were grown up to ∼10^6^ in a 5% CO_2_ atmosphere and were then infected with 11168H_CBA_, 11168H_AC/CBA_ or 11168H_AP/CBA_ *C. jejuni* at MOI of 200:1 for 3□hrs as described previously [13]. The monolayers were then washed three times with PBS, incubated in DMEM containing gentamicin (100□µg/mL) for 2□h at 37□°C to kill extracellular bacteria, the cells were then washed 3x with PBS and then lysed with 0.1% (v/v) Triton X-100. The cell lysates were serially diluted and plated onto blood agar plates and incubated for 48□h before colonies were enumerated. Experiments were performed in triplicates of three biological replicates. To normalize the numbers of intracellular bacteria recovered, the initial bacterial inoculum was always plated.

### *Galleria mellonella* infection model

*Galleria mellonella* larvae (LiveFoods) were kept on wood chips at room temperature. Experiments were performed as previously described [14]. Briefly, 11168H_CBA_, 11168H_AS/CBA_ or 11168H_AP/CBA_ *C. jejuni* were suspended in PBS to give OD_600nm_ 0.1 and 10□µl volume of this suspension (∼10^6^) was injected into the right foremost leg of the *G*.□*mellonella* larvae by microinjection (Hamilton) and incubated at 37°C. For each strain, 10 larvae with similar weight were used per replicate. Mortality was observed at 24 hrs intervals for 72 hrs.

### Statistical analysis

All experiments presented are at least three biological replicates. All data were analyzed using Prism statistical software (Version 9, GraphPad Software). Values were presented as standard deviation and variables were compared for significance using student t-tests to obtain *p*-values unless otherwise stated.

## Results and Discussion

### Survival of *C. jejuni* in Acanthamoeba increases subsequent invasion

*C. jejuni* cells that had survived within *Acanthamoeba spp*. were tested for their ability to invade T84, Caco-2, *A. castellanii* and *A. polyphaga* cells **Fig. 2**. Invasion of all cells (human cell lines and amoebae cells) increased significantly (*p*<0.05) with bacteria harvested from amoebae, 11168H_AC/CBA_ or 11168H_AP/CBA_, more than with 11168H_CBA_ *C. jejuni*.

**Figure 2.**
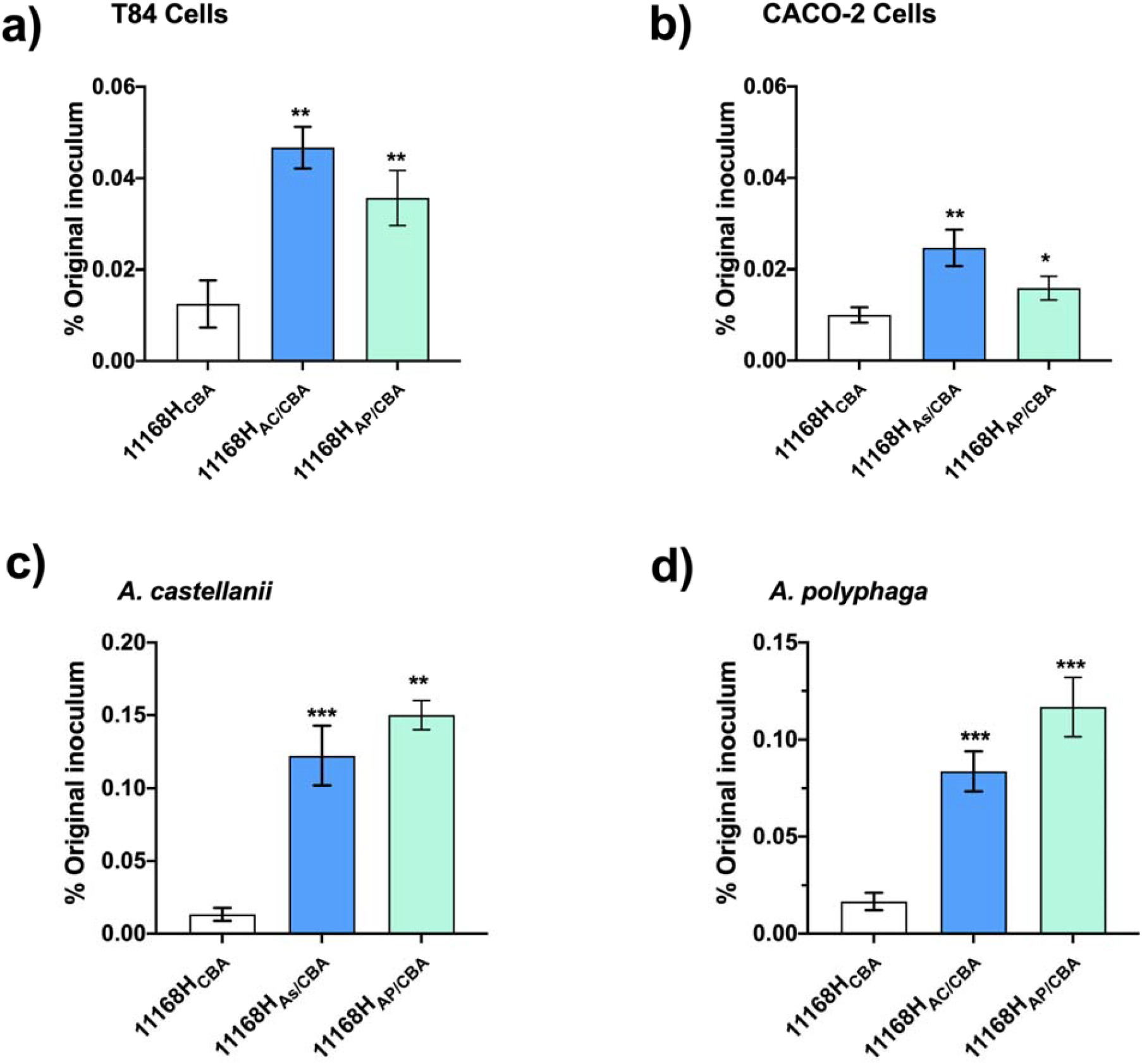
*C. jejuni* 11168H harvested from *Acanthamoeba spp*. **a)** invasion of T84 cells; **b)** Caco-2 cells; **c)** *A. castellanii*; and **d)** *A. polyphaga*. Invasion of strain 11168H_CBA_, 11168H_AC/CBA_ or 11168H_AP/CBA_ were determined by infection of the cell lines for 3 hrs and enumerated after 100 μg/mL gentamicin treatment and lysis of cell. Data is presented as % of the original inoculum. Error bars represent SD from three independent experiments. *p≤ 0.05, **p ≤ 0.01, ***p ≤ 0.001.

We observed that bacteria passaged through amoebae had an increased capacity to invade human epithelial cells compared to non-passaged 11168H_CBA_ *C. jejuni* with a relative increase in invasion of ∼3.7-fold for 1168H_AC/CBA_ and ∼2.8-fold for 11168H_AP/CBA_ in T84-cells **(Fig. 2a)**. In Caco-2 cells, we also observed a significant increase in relative invasion, although to a lesser extent, with ∼2.4-fold for 11168H_AC/CBA_ and ∼1.6-fold for 11168H_AP/CBA_ **(Fig. 2b)**. The levels of invasion varied dependent on the cell type, however, it was always significantly higher than that of 11168H_CBA_ bacteria. These results indicated that *C. jejuni* cells that survived intracellularly in amoebae undergo priming and are more invasive, this increased invasiveness could lead to a more severe disease in humans. This priming adaptive response has been reported in bacteria [15], including *C. jejuni* adaptive tolerance to low pH [16]. To our knowledge, this is the first study that shows *C. jejuni* survival within amoebae enhances subsequent invasion of human epithelial cells. This provides novel insight into the interactions of *C. jejuni* with protists.

To determine whether this increase in invasion would also be observed in amoebae, we performed re-infection of the *Acanthamoeba spp*. with 11168H_CBA_, 11168H_AC/CBA_ or 11168H_AP/CBA_ *C. jejuni*. Interestingly, re-infection of *Acanthamoeba spp*. showed a significant increase in invasion with bacteria passaged through amoebae compared to non-passaged bacteria, 11168H_CBA_. *A. castellanii* invasion showed significant increase of ∼9.2-fold with 11168H_AC/CBA_ and ∼11-fold with 11168H_AP/CBA_ *C. jejuni* **(Fig. 2c)**. The same trend was observed for *A. polyphaga* re-infection, ∼5.0-fold increase in invasion with 11168H_AC/CBA_ and ∼7.0-fold with 11168H_AP/CBA_ *C. jejuni* **(Fig. 2d)**. To ensure that the observed increased in invasion was not caused by resistance to gentamicin; sensitivity tests were performed, and revealed no significant (p<0.05) differences between 11168H_CBA_ and amoebae recovered *C. jejuni* (data not shown).

The dramatic invasion of 11168H_AC/CBA_ and 11168H_AP/CBA_ within amoebae may be attributed to the higher background of non-specific uptake of bacteria by amoebae including non-pathogenic bacteria. However, given that increase in invasion was observed across all the cell types used in this study, it is more likely that amoebae recovered bacteria have pre-adapted to subsequently invade and survive at a greater rate than that of 11168H_CBA_ bacteria. This finding may be unsurprising, since the symbiotic relationship between amoebae and bacteria has been thought to pre-adapt intracellular microorganisms to survive in other cells including human macrophages [17]. This eco-evo hypothesis [18] is observed with *Chlamydia spp*. and *Legionella pneumophilia* which use similar strategies to interact with various hosts cells and most probably evolved over millions of years during bacterial interactions with primitive unicellular eukaryotes [17, 19, 20]. This greater invasiveness of *C. jejuni* within amoebae could facilitate longer survival which may lead to an increased ability of *Campylobacter* to survive and subsequently transmit to humans from environmental sources.

### *Acanthamoeba spp*. are a transient host for *C. jejuni*

There have been conflicting reports on intracellular multiplication of *C. jejuni* within amoebae [5, 21]. Whilst some have reported *C. jejuni* is capable of multiplying within *Acanthamoeba spp*. [3, 12, 21] others have been unable to observe intracellular multiplication [4, 22]. In our model, we were unable to detect intra-amoebae multiplication of 11168H and 81-176 *C. jejuni* strains **(Fig. 3)** in different media, PYG **(Fig. 3a and b)** or in brucella broth (a highly nutritious media used to enrich *C. jejuni* growth and can sustain *Acanthamoeba spp*.) **(Fig. 3c and d)**, aerobically at 37 °C as previously described [12, 21]. We found that *C. jejuni* cells were undetected at 72 hrs post-infection. These findings led us to conclude that amoebae, at least, the *Acanthamoeba spp*. used in this study are a transient host for *C. jejuni*. A previous study presented a hypothetical model suggesting that in the environment *C. jejuni* multiplies within amoebae, potentially bursting out and re-invading neighboring amoebic cells [12]. The experimental data presented here partly confirms their hypothesis, however, since we did not observe intracellular multiplication of *C. jejuni*, we propose that it is more likely that in the environment outside the host, *C jejuni* would invade *Acanthamoeba* cells briefly, but frequently, with increased invasion efficiency. This strategy increases the chances of this bacterium to be transmitted to warm blooded avian and mammalian hosts.

**Figure 3.**
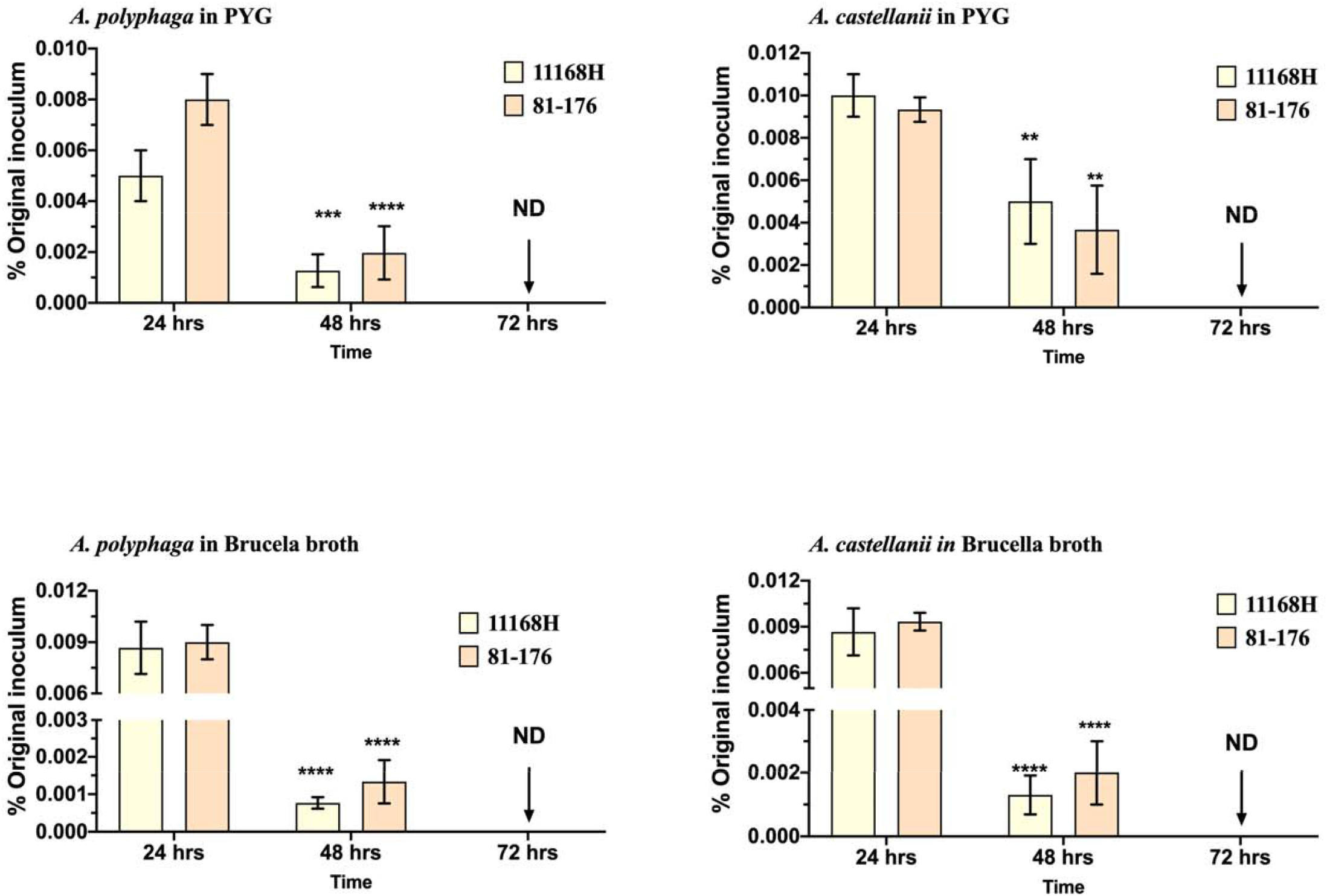
Long-term survival of *C. jejuni* with *Acanthamoeba spp*. *C. jejuni* strains 11168H or 81-176 survival in **a)** *A. polyphaga*; **b)** *A. castellanii* in PYG media, and **c)** *A. polyphaga*; **d)** *A. castellanii* in brucella broth at 37 °C in aerobic conditions. Amoebae were lysed for enumeration of live bacteria at 24 hrs interval for 72 hrs after 10 μg/mL gentamicin treatment. Data is presented as percentage of the original inoculum. Error bars represent SD from three independent experiments. Two-way ANOVA multiple comparison was used to test for significance; **p ≤ 0.01, ***p ≤ 0.001, ****p≤ 0.0001. ND = no bacteria detected.

### *C. jejuni* survival in amoebae is not cytotoxic to *Galleria mellonella* larvae infection model

Increased invasion of the different cell lines prompted us to examine whether *C. jejuni* passaged through amoebae would be more cytotoxic for *G. mellonella* larvae compared to 11168H_CBA_ bacteria. Using this surrogate infection model, we did not observe any significant cytotoxic differences between 11168H_CBA_, 11168H_AC/CBA_ or 11168H_AP/CBA_ *C. jejuni* towards *G. mellonella* larvae **Fig. 4**.

**Figure 4.**
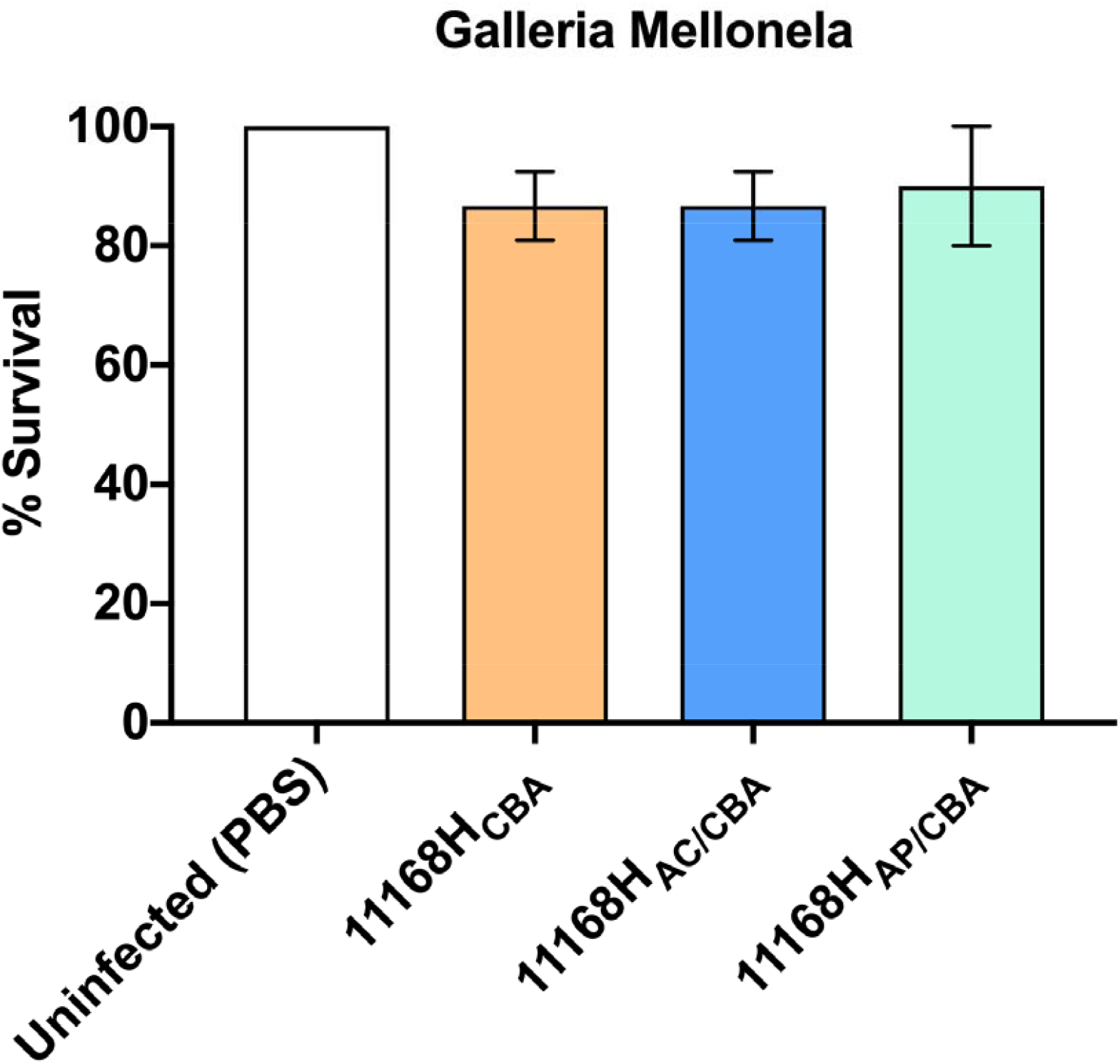
The effect of 11168H_CBA_, 11168H_AC/CBA_ and 11168H_AP/CBA_ in the *Galleria mellonella* infection model. *G. mellonella* larvae were injected with a 10 μl inoculum of *C. jejuni* 10^6^ CFU by microinjection in the right foremost leg. Larvae were incubated at 37°C, with survival and appearance recorded after 72 hrs. PBS injection control was used. For each experiment, 10 *G. mellonella* larvae were infected, and experiments were repeated in triplicate. Error bars represent SD. Cytotoxicity was monitored at 24 hrs intervals for 72 hrs.

Although this infection model has been previously used to determine *C. jejuni* cytotoxicity [13, 14, 23], we cannot rule out that significant differences may be observed in avian and mammalian host cells. A previous study by Snelling *et al*., showed increased chicken colonization after 7-days post infection with intra-amoebae *C. jejuni* [24]. It would be worth studying cytotoxicity in chicken colonization/infection models.

### *A. castellanii* supports greater survival of *C. jejuni*

In the environment, *C. jejuni* would encounter multiple species of amoebae. We examined the invasiveness and survivability of six *C. jejuni* strains; 11168H, 81-176, 12912, M1, 81116 and RM1221 within *A. castellanii* and *A. polyphaga* **Fig. 5**. These strains were selected because of their diverse genetic backgrounds and sources, thus our observations are more representative of the *C. jejuni* species **(Table 1)**.

**Figure 5.**
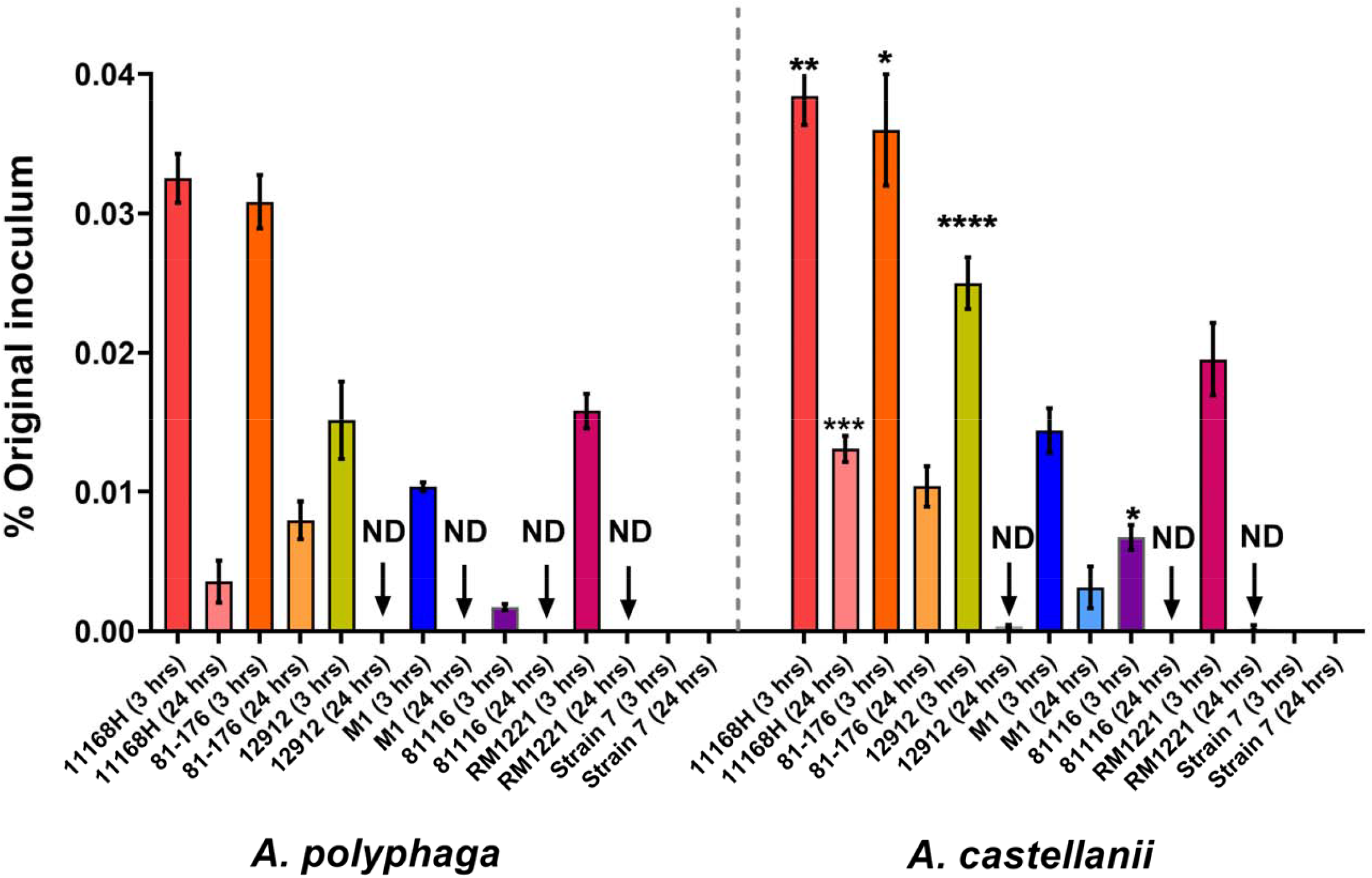
Quantification of *C. jejuni* strains survival within *Acanthamoeba species*. *C. jejuni* strains 11168H, 81-176, 12912, M1, 81116 and RM1221 in *A. polyphaga* and *A. castellanii*. Quantification of intracellular bacteria was determined by viable counts at 3 hrs and 24 hrs after amoebae infection at 25 °C in aerobic conditions. Data is presented as percentage of the original inoculum. Error bars represent SD from three independent experiments. Two-way ANOVA multiple comparison was used to test for significance; *p≤ 0.05, **p ≤ 0.01, ***p ≤ 0.001, ****p≤ 0.0001. ND = no bacteria detected.

We observed a general trend of a greater survival rate of *C. jejuni* strains within *A. castellanii* compared to *A. polyphaga* in PYG at 25 °C under aerobic conditions. Our results show differences in invasiveness and survival capabilities between the range of the *C. jejuni* strains tested. Interestingly, these results are similar to previous studies that correlated invasiveness and survivability of *C. jejuni* is both bacterial strain and host cell-dependent [25-28]. In our amoebae model, the greater survivability of *C. jejuni* within *A. castellanii* seems to be a host susceptibility factor rather than being bacterial induced. This phenomenon was also reported in other bacteria, where greater levels of invasion and intracellular survivability of *Listeria monocytogenes* was observed in *A. castellanii* [29] compared to other *Acanthamoeba spp*.. A recent review proposed that invasion and intracellular occurrences of microbes within amoebae is dependent on the genotype of the host [30]. Based on 18S RNA sequence studies, *A. castellanii* is from the T4 genotype [31] whilst *A. polyphaga* is from the T2 genotype [32], although how host genotype plays a role in intracellular survival remains unknown.

The mechanisms of survival within amoebae has been compared to that of macrophages, and a review by Vieira *et al*., [5] predicted various factors that *C. jejuni* could utilize to invade and survive within amoebae cells. Although survivability of *C. jejuni* in our model is most likely host-dependent, it would be intriguing to determine the bacterial factors involved. Unlike other enteric pathogens, the *C. jejuni* genome is relatively small at 1.6 Mb [33, 34], and this bacterium would plausibly use the same factors to invade and survive within amoebae as it would for avian and mammalian host cells. Therefore, elucidating these factors may give new insights into *Campylobacter* infection which compared to other enteric pathogens is poorly understood. Future studies could use molecular tools such as genome-wide transposon mutant libraries of multiple *C. jejuni* strains like those used by de Vries *et al*., [35] to help improve our understanding of *Campylobacter* infection.

In conclusion, given the two species of amoebae and diverse selection of *C. jejuni* strains tested, this consistent data provides insight into a natural phenomenon that will be important in *Campylobacter* survival, transmission and infection. We propose that *Acanthamoeba spp*. are a transient host for *C. jejuni* and survival within this “Trojan horse” environment subsequently increases *C. jejuni* invasiveness.

## Acknowledgements

The authors would like to thank Claire Rogers and Debbie Nolder for their advice on handling *Acanthamoeba*. Geunhye Hong for seeding mammalian epithelial cells line and Elizabeth McCarthy for generating microscopy images. FN conceptualized the study, designed experiments and performed all experiments; FN and BW wrote the manuscript. This work was supported by Biotechnology and Biological Sciences Research Council Institute Strategic Program BB/R012504/1 constituent project BBS/E/F/000PR10349 to B.W. All authors declare they have no known conflict of interest.

